# Sodium-mediated plateau potentials in an identified decisional neuron contribute to feeding-related motor pattern genesis in *Aplysia*

**DOI:** 10.1101/2023.03.30.534938

**Authors:** Alexis Bédécarrats, John Simmers, Romuald Nargeot

## Abstract

Motivated behaviors such as feeding depend on the functional properties of decision neurons which provide the flexibility required for behavioral adaptation. Here, we analyzed the ionic basis of the endogenous membrane properties of an identified decision neuron (B63) that drive radula biting cycles underlying food-seeking behavior in *Aplysia*. Each spontaneous bite cycle arises from the irregular triggering of a plateau-like potential and resultant bursting by rhythmic subthreshold oscillations in B63’s membrane potential. In isolated buccal ganglion preparations, and after synaptic isolation, the expression of B63’s plateau potentials persisted after removal of extracellular calcium, but was completely suppressed in a TTX-containing bath solution, thereby indicating the contribution of a transmembrane Na^+^ influx. Potassium outward efflux through TEA- and calcium-sensitive channels was found to contribute to each plateau’s active termination. This intrinsic plateauing capability, in contrast to B63’s membrane potential oscillation, was blocked by the *I*_CAN_ blocker flufenamic acid (FFA). Conversely, the SERCA blocker cyclopianozic acid (CPA), which abolished the neuron’s oscillation, did not prevent the expression of experimentally evoked-plateau potentials. These results therefore indicate that the dynamic properties of the decision neuron B63 rely on two distinct mechanisms involving different sub-populations of ionic conductances.

**NEW & NOTEWORTHY:** Here, we report an endogenous plateau property underlying bursting in a bilateral pair of buccal ganglion pacemaker neurons (B63) which trigger individual motor pattern cycles for food-seeking behavior in the marine mollusk *Aplysia*. The ionic mechanisms of this membrane bistability in B63 rely critically on voltage-dependent sodium inward currents. The expression of these plateau properties and the underlying endogenous oscillatory pacemaker drive can be dissociated pharmacologically, indicating that the two intrinsic properties depend on different sets of conductances. Our results thus shed new light on the mechanisms of spontaneous decision making in *Aplysia*.

## INTRODUCTION

Bistable membrane behavior giving rise to depolarized, burst-generating plateau potentials is a widespread endogenous membrane property of neurons in central pattern generating networks for diverse rhythmic behaviors such as breathing, chewing or walking (Russell and Hartline 1978; Hounsgaard et al., 1984; Kiehn, 1991; Onimaru et al., 1996; Russo and Hounsgaard, 1996; Brocard et al. 2006; Husch et al. 2015; Boeri et al., 2018). Although many neurons are capable of action potential bursting, the contribution of an underlying plateau potential capability has not always been established.

Food-seeking behavior in *Aplysia* partly consists of cycles of protraction/retraction of the tongue-like radula which are triggered by decision neurons located in the buccal ganglia (Cropper et al. 2004; Nargeot and Simmers, 2011). The decision-making process responsible for the expression of radula motor pattern cycles is based on intracellular calcium release-derived oscillations in membrane potential of two bilateral pacemaker neurons, B63, and their excitatory synaptic transmission to electrically-coupled buccal ganglion neurons, including the B31/B32 motor neurons (Bédécarrats et al., 2021; Susswein et al., 2002). In contrast, earlier experimental and computational data suggested that the B63 depolarizations underlying the impulse bursts that drive individual radula bite cycles may not rely on an intrinsic property of this neuron *per se*. Rather, they were proposed to result from B63’s fast cholinergic synaptic excitation of the buccal network’s B31/B32 neurons and a prolongation of the latters’ depolarizing response and associated firing by voltage-activated autapses. This in turn provides depolarizing-sustaining feedback excitation of B63 through electrical coupling (Susswein et al. 2002; Cataldo et al. 2006; Costa et al., 2020; Momohara et al. 2022). However, we recently found that B63 can express spontaneous or experimentally-evoked plateau potentials in modified bathing saline that blocks chemical synapses in the buccal network (Bédécarrats et al, 2021). In the present study we now provide evidence for the ability of B63 to produce plateau potentials independently of B31/B32 depolarization under standard physiological conditions. We then characterize the ionic basis of these B63 plateau potentials under complete chemical synapse blockade. Our results therefore suggest that in addition to the previously described synaptic mechanism for plateau-like behavior, the B63 pacemaker neurons play an active role in the decision-making process not only through their endogenous oscillatory property, but also via a distinct intrinsic plateauing capability.

## METHODS

### Animals

*Aplysia californica* purchased from the University of Florida were housed in tanks containing fresh aerated sea water (~15°C) and fed with seaweed (*Ulva lactuca*) obtained from the Station Biologique at Roscoff, France.

### Isolated buccal ganglia preparations

Animals were anesthetized with an injection of a 60 ml isotonic MgCl_2_ solution (in mM: 360 MgCl_2_, 10 HEPES, adjusted to pH 7.5). Buccal ganglia were dissected under artificial sea water (ASW, in mM: 450 NaCl, 10 KCl, 30 MgCl_2_, 20 MgSO_4_, 10 CaCl_2_, 10 HEPES, adjusted to pH 7.5). The isolated ganglion preparations were maintained at 15°C with a Peltier cooling device and bathed in standard ASW or modified saline during subsequent pharmacological experiments.

### Modified saline & pharmacology

A ‘Low Ca+Co’ solution was used to block all network chemical synaptic connections and was composed of: (in mM) 446 NaCl, 10 KCl, 30 MgCl_2_, 20 MgSO_4_, 3 CaCl_2_, 10 CoCl_2_, 10 HEPES, adjusted to pH 7.5. NaCl concentration was adjusted to maintain the same osmolarity as ASW. Electrophysiological recordings under this saline started at least 20 min after perfusion onset, which were required to completely block chemical synapses and allow for recovery of neuronal resting membrane potentials to at least −50 mV (Bédécarrats et al., 2021). A calcium-free solution (‘0 Ca+Co’) in which all calcium (CaCl_2_) was replaced with equimolar (10 mM) cobalt (CoCl_2_) was composed of (in mM): 450 NaCl, 10 KCl, 30 MgCl_2_, 20 MgSO_4_, 10 CoCl_2_, 10 HEPES, 0.5 EGTA, adjusted to pH 7.5). D-tubocurarin (Merck-Sigma-Aldrich) was used at a concentration of 1 mM in ASW. Tetrodotoxin (TTX, Tocris) was diluted in distilled water from a 0.1 mM stock solution and added to Low Ca+Co saline at a concentration of 1 μM. Tetraethylammonium (TEA, Merck-Sigma-Aldrich) was diluted in Low Ca+Co saline at a concentration of 50 mM. Flufenamic acid (FFA, Merck-Sigma-Aldrich) was first diluted in 100 mM DMSO and then in Low Ca+Co saline at a concentration of 100 μM. Cyclopianozic acid (CPA, Merck-Sigma-Aldrich) was diluted to 20 mM in DMSO and used at a final concentration of 20 μM in Low Ca+Co.

### Electrophysiological recordings

Spontaneously expressed cycles of radula motor output (buccal motor patterns, BMPs) were monitored with wire pin extracellular electrodes placed in direct contact with the cut stumps of the protraction I2 (I2 n.), retraction 2,1 (n. 2,1) and radular closure motor (Rn.) nerves. Intracellular electrodes were made from pulled glass capillaries (15–30 MΩ) filled with KCH_3_CO_2_ (2 M). Intracellular signals were amplified with an Axoclamp-2B amplifier (Molecular Devices, Palo Alto, CA), digitized with a CED (Cambridge Electronic Design) interface (sampling rate 5 kHz), then acquired and analyzed with Spike 2 software (Cambridge Electronic Design, UK). B63 interneurons and B31/B32 motoneurons were identified as previously described (Susswein and Byrne, 1988; Nargeot et al. 2009). Relatively brief intracellular current pulses (duration 5 s or 8 s) were injected to trigger plateau-like activity in B63 under ASW. Longer pulses of 30 s or 60 s were used to trigger plateaus under chemical synapse blockade in Low Ca+Co saline.

### Data analysis

The amplitudes of B63’s spontaneous plateau-like potentials were measured from the membrane potential value at the base of the first action potential until the maximum level of step-like depolarization. The simultaneous voltage variation in a contralateral B31/B32 neuron, which is postsynaptic to a given B63, was also measured. The amplitudes of evoked plateau-like potentials in each recorded B63 and in the contralateral B31/B32 were measured from the resting potential immediately prior to current pulse injection to the maximum level of pulse-triggered depolarization.

Underlying subthreshold oscillations in membrane potential of B63 neurons were analyzed between a bandwidth of 0.00390 to 0.125 Hz by Fast Fourier Transform (FFT) analysis using R software (R Development Core Team, 2019). Intracellular voltage recordings were first smoothed with a Spike 2 ‘Smooth’ filter with a time constant of 500 ms to suppress action potentials and down sampled at 1 Hz to decrease the time of computer processing. The peak magnitudes of spectral densities were determined for 600 s excerpts of B63 intracellular recordings. Sinusoidal waveforms corresponding to plateau-triggering oscillations detected on the FFT periodograms were computed and reconstructed from Wavelet decomposition using the R-CRAN package ‘WaveletComp’ (Roesch and Schmidbauer, 2018).

### Statistical analyses

Two-tailed Mann-Whitney *U* tests were used for unpaired comparisons of two independent groups. Comparisons of the proportions of neurons from different preparations expressing evoked plateau-like potentials were made using the two-tailed Fisher’s exact test in unpaired-sample procedures. A Kruskal–Wallis test (X^2^) was used to assess the significance of the overall group differences between three independent groups. Post-hoc pairwise comparisons were performed with a Dunn’s test (q). All statistical tests were conducted with Prism 9 software (GraphPad) and any comparison differences were considered significant for a probability level of p < 0.05. Bars in figures represent Mean ± SEM, \**p* < 0.05, \*\**p* < 0.01, \*\*\**p* < 0.001, n.s., non significant.

## RESULTS

### Plateau potentials in B63 can occur independently of B31/B32 plateauing

The premotor pacemaker B63 neuron spontaneously expresses an endogenous oscillation of its membrane potential (Figure 1A-C) that occasionally triggers a large amplitude, plateau-like depolarization and intense burst discharge associated with buccal motor pattern (BMP) genesis (Figure 1A). This activity in the bilateral pair of B63 neurons was previously thought to drive BMP generation through a combination of monosynaptic fast cholinergic EPSP production in, and electrical coupling with, the contralateral motor neurons B31/B32 (Hurwitz et al. 1997; Hurwitz et al. 2003). In this scheme, when sufficiently depolarized, the B31/B32 cells express a plateau-like state and a prolonged burst of action potentials via self-excitatory synapses, which in turn sustains B63’s depolarization through their electrical coupling (Figure 2A1,2) (Saada et al., 2009).

**Figure 1.**
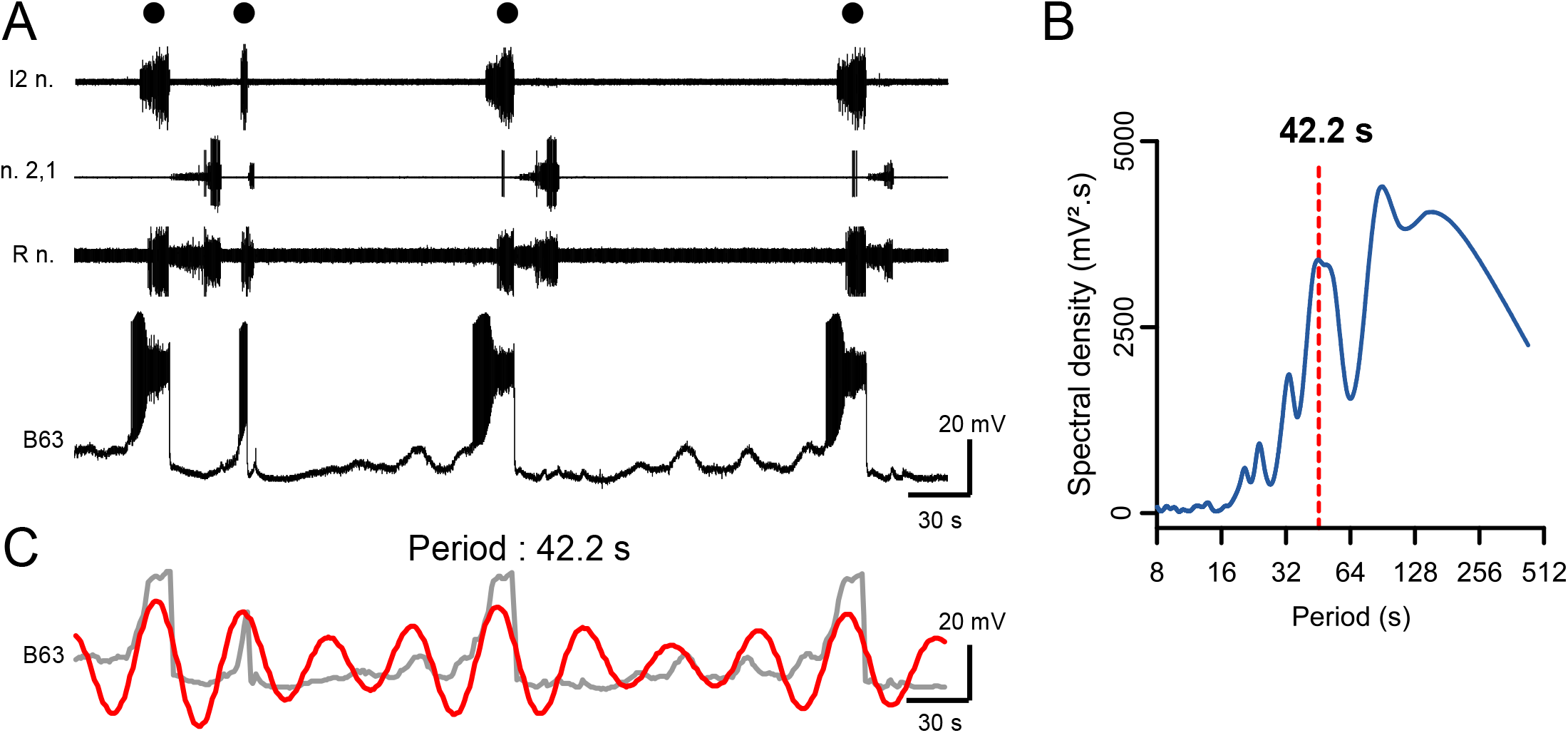
Spontaneous expression of B63 neuron plateau-like potentials and underlying rhythmic membrane potential oscillations associated with buccal motor pattern (BMP) genesis. **A.** Simultaneous extracellular recordings from motor nerves of isolated buccal ganglia (top 3 traces) and an intracellular recording of a B63 interneuron (bottom trace). Black dots indicate expression of a spontaneous BMP associated with each large amplitude, prolonged depolarization and burst of action potentials in the B63. **B.** FFT periodogram of the B63 recording as illustrated in A, indicating a rhythmic underlying fluctuation in the neuron’s membrane potential (dashed line, period 42.2 s). **C.** Wavelet-based reconstruction of this fastest (period 42.2 s) voltage oscillation (red trace) superimposed on the smoothed B63 membrane voltage trace (grey). Each plateau-like potential arises occasionally from the positive peak of an oscillation cycle. Buccal nerve abbreviations: l2n., n.2,1, Rn., are protractor, retractor and closure motor nerves respectively.

**Figure 2.**
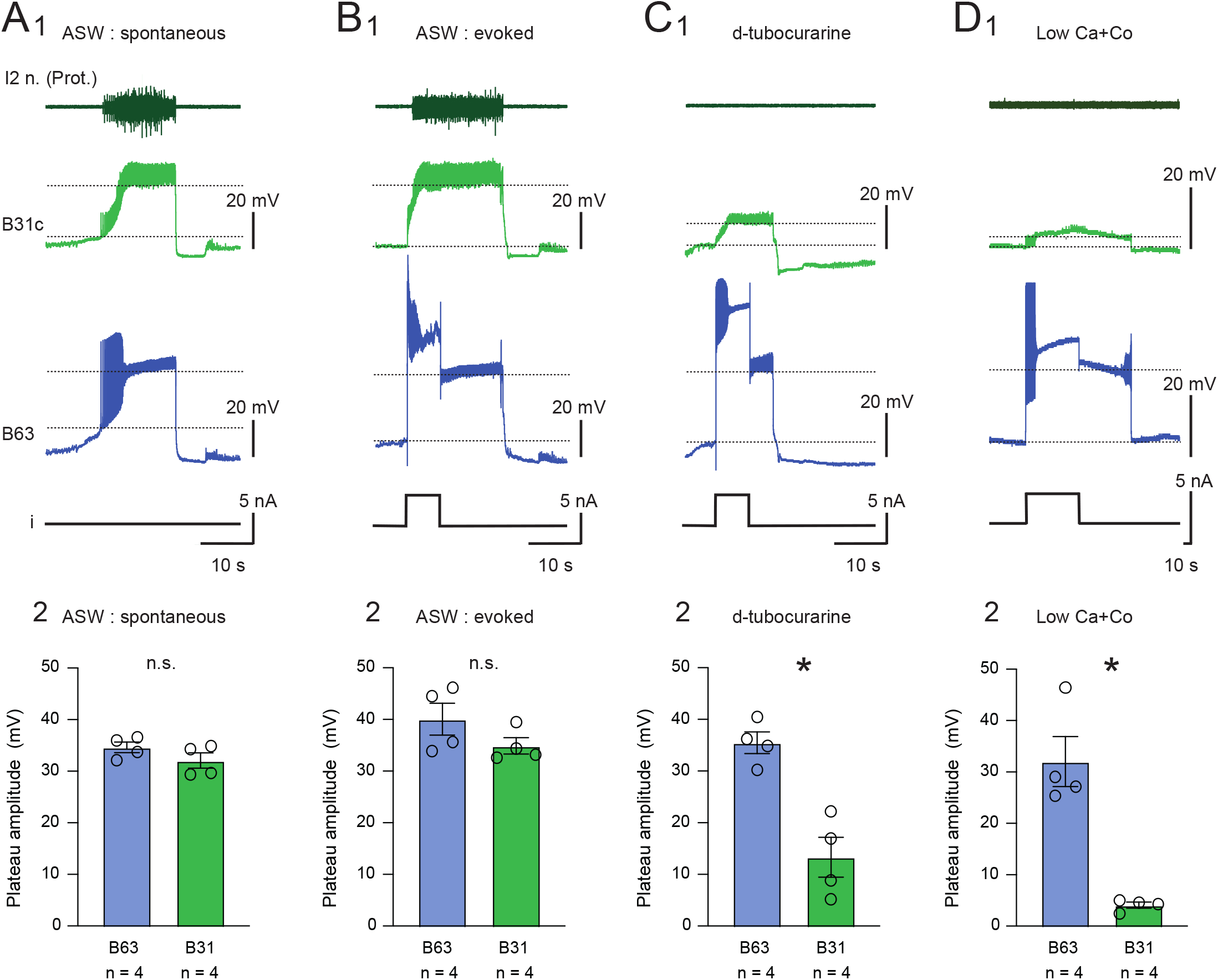
Endogenous plateau property of the B63 neuron. Representative intracellular recordings of spontaneous and evoked activity in B63 neurons and a contralateral protractor motoneuron B31c (A1-C1 are from a same preparation). In each case, a simultaneous extracellular recording of the protraction nerve I2 n. (upper traces) and a monitor of current injected into B63 (i, bottom traces) are also shown. Dotted lines indicate baseline and maximum membrane potential levels used for determining B63 and B31 depolarizing response amplitudes. **A1.** Spontaneous plateau-like potential expression in a B63 and B31 cell under standard ASW. Note the resulting impulse burst occurring in the axons of B31/32 motor neurons conveyed in l2n. **A2.** Group data: the maximal amplitude of the membrane depolarizations underlying burst discharge were not significantly different in B63 (34.63 ± 1.025 mV) and B31 (32.07 ± 1.477 mV; *U* = 4, p = 0.343). **B1.** Plateau-like potentials evoked in the same B63 and a B31neurons by depolarizing current pulse injection into the former under standard ASW. **B2.** No significant difference was evident between the mean amplitudes of plateaus expressed in B63 (40.05 ± 3.082 mV) and B31/B32 neurons (34.90 ± 1.587 mV; *U* = 3, p = 0.200). **C1.** A plateau potential evoked in the current pulse-injected B63 under 1 mM d-tubocurarine. **C2.** D-tubocurarine significantly reduced the amplitudes of B31’s depolarizing responses (13.34 ± 3.842 mV) as compared to those of B63 (35.47 ± 2.088 mV; *U* = 3, p = 0.200). Note the absence of B31/32 axonal discharge in I2n. **D1.** Evoked B63 plateaus under complete synaptic isolation in Low Ca+Co saline. **D2.** Blockade of the buccal network chemical synapses caused a significant further decrease in the depolarizing responses of B31/B32 (4.1 ± 0.575 mV) compared to the plateaus evoked directly in B63 (32.04 ± 4.863 mV; *U* = 0; p = 0.023).

In a first set of experiments using isolated preparations of buccal ganglia under normal saline (ASW), we asked whether the plateau-like depolarizations in B63 can be expressed in the absence of corresponding B31/B32 neuron activation. With sufficient strength, brief pulses (5 s or 8 s) of depolarizing current injected into a recorded B63 neuron could trigger long-lasting plateau-like depolarizations of similar amplitude in both the injected B63 itself and a simultaneously recorded B31 neuron (Figure 2B1,2). Since this latter neuron’s depolarization is thought to be triggered by fast cholinergic excitation from B63, the nicotinic antagonist D-tubocurarin (1 mM) was applied to test whether the two neuronal responses could be dissociated. The presence of D-tubocurarin in the bath saline effectively blocked fast EPSP production in B31/B32 by B63. Consequently, the formers’ depolarization in response to B63 activation was now strongly reduced (Figure C1,2) and was insufficient to reach threshold for impulse firing, as indicated by the total absence of spikes in B31/32 axons monitored in nerve I2 (Figure C1). In contrast, the initial injected current pulse still elicited a plateau-like depolarization and prolonged burst firing in B63 itself, both of which outlasted the triggering pulse and with a plateau amplitude that remained unchanged from control (compare Figure 2C, 2B).

This dissociation of the membrane behavior of the B63 and B31/32 neurons was further confirmed by exposure to a Low Ca+Co bath solution to block chemical synapses throughout the buccal network (Figure 2D1, 2). In this condition, brief current injection into a B63 again elicited a sustained, large amplitude voltage shift but produced only a weak subthreshold depolarization in a simultaneously recorded B31 neuron. Presumably, this residual response resulted from the latter’s electrical coupling with B63. Altogether, these results are consistent with the idea that although chemical excitation from B63 acting in combination with B31/B32’s autapses may contribute to plateau-like behavior in these two cell types, this synaptic mechanism is not exclusively responsible for B63’s plateauing capability, but rather, it results at least in part from an endogenous membrane property.

### Ionic mechanisms for B63 plateauing

Plateau potential generation is the expression of a bistable, all-or-none electrical behavior that allows cells to maintain a lasting depolarized membrane potential before switching back to resting potential, either spontaneously or in response to hyperpolarizing synaptic inputs or negative current injection (Russell and Hartline, 1982; Hartline and Graubard, 1992). The depolarized plateau state of this bistability is typically maintained by persistent inward currents through voltage-dependent cationic channels, while plateau termination is due to voltage-induced closure of these channels and/or to outward currents though calcium-gated potassium channels (Golowasch and Marder, 1992; Russo and Hounsgaard, 1996; Tabak et al., 2011). To investigate the contribution of Na and/or Ca^2+^ ions to the onset and maintenance of the active depolarized state of the B63 neurons, experiments (n = 4) were conducted in which the sodium channel blocker TTX (50 μM) was added to the Low Ca+Co experimental solution. In all preparations under this condition, injected depolarizing current pulses failed to trigger B63 plateau potentials (Figure 3A, C). In contrast, in a separate group of preparations (n = 4) that were bathed in Ca^2+^ free saline to which the calcium channel blocker Co^2+^ had been added, B63 was still able to express a sustained plateau potential following positive current pulse injection, and then terminated spontaneously or in response to a brief hyperpolarizing current pulse (Figure 3B, C). On this basis, therefore, extracellular Na^+^ influx through TTX-sensitive membrane channels, but not extracellular Ca^2+^ influx, appears to be necessary for the expression of B63’s endogenous plateau property.

**Figure 3.**
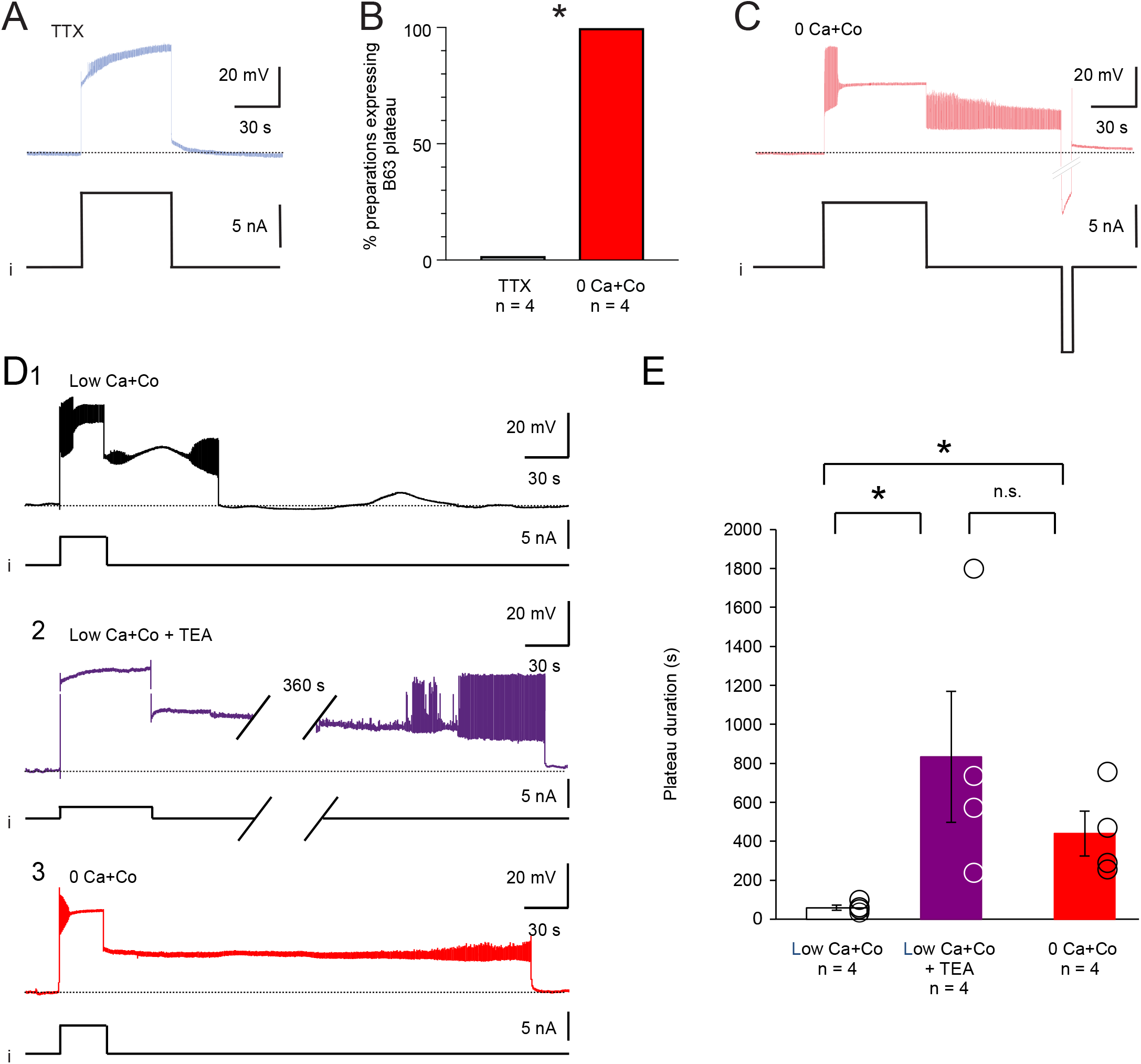
Ion conductances involved in the maintenance and termination of B63 plateau potentials. **A.** TTX (50 μM) added to Low Ca+Co bathing saline prevented the ability of positive current pulse injection (I, lower trace) to elicit a prolonged B63 plateau potential. **B.** Percentage comparison of preparations that successfully expressed evoked B63 plateaus under 0 Ca^2+^ or Low Ca+Co saline containing 50 μM TTX (p = 0.029). **C.** Under 0 Ca^2+^ saline, a plateau potential could still be elicited in a B63 neuron by a positive current pulse and terminated with an injected negative pulse (see i). **D.** Timing of spontaneous B63 plateau termination under different saline and pharmacological conditions. Plateau termination in Low Ca+Co saline (D1) was considerably delayed in the presence of 50 mM TEA (D2, breaks in the recording traces correspond to 360 s) or in the complete absence of saline Ca^2+^ (D3). **E.** The plateau duration was significantly different between groups (X^2^_(2)_ = 7.539, p = 0.023), being longer in the presence of TEA in Low Ca+Co (q = −2.549, p = 0.016) or after the removal of calcium (0 Ca+Co; q = −2.157, p = 0.046) than in the Low Ca+Co solution alone. No significant difference was evident between the 0 Ca+Co and TEA groups (q = −0.392, p = 1.000).

In normal saline (ASW) conditions, the spontaneous termination of individual plateaus and associated impulse bursts in B63, as in other buccal neurons generating the protractor phase of each BMP, is triggered by an inhibitory synaptic drive from neurons active during the retraction phase of each radula bite cycle (Hurwitz and Susswein, 1996; Sasaki et al. 2013). Under Low Ca+Co or 0 Ca+Co solutions, which suppress these inhibitory influences, B63 expressed greatly extended plateau potentials that eventually terminated spontaneously (Figure 3D1, D3, E). This step-change cessation could also be prematurely induced by the intracellular injection of a hyperpolarizing current (see Figure 3C), thereby suggesting the contribution of a voltage-dependent mechanism.

To test for a probable contribution of potassium channels in B63’s plateau termination, preparations were superfused with or without the potassium channel blocker TEA (50 mM) in Low Ca+Co saline. In the presence of TEA (n = 4), as compared to the blocker’s absence (n = 4), the durations of B63’s plateau potentials were considerably and significantly prolonged (Figure 3D2, compare with D1, E). Plateau durations were also considerably modified by changes in the concentration of calcium ions, with plateaus lasting several tens of seconds in the presence of Ca^2+^ (3 mM in the Low Ca+Co solution, n = 4), but were maintained for hundreds of seconds in the cation’s absence (0 Ca+Co solution, n = 4) (Figure 3D3, compare with D1, E). These findings thus lead to the conclusion that a voltage-dependent potassium current regulated by calcium contribute to B63’s plateau termination.

### Different cation channels underlie B63’s endogenous plateau and oscillatory properties

In normal ASW conditions, plateau potentials in the B63 neuron are triggered spontaneously by an ongoing low amplitude oscillation in the interneuron’s membrane potential (see Figure 1) that arises from organelle-derived fluxes in intracellular calcium (Bédécarrats et al., 2021). Both the plateaus (as described above) and their underlying voltage oscillation are blocked by TTX, indicating in each case, a contribution of sodium and/or non-selective cation membrane channels (see Figure 3A1 and Bédécarrats et al., 2021). To determine whether the same or different membrane channels are implicated in the expression of these endogenous properties, the specific effects of two different pharmacological agents was tested. Firstly, the membrane potential oscillation of B63 was suppressed by the presence of reticulum endoplasmic calcium pump inhibitor CPA (20 μM in a Low Ca+Co solution; Figure 4A1, C; see also Bédécarrats et al., 2021), and consequently, also prevented the spontaneous expression of plateau potentials (n = 8; Figure 4A1). Nonetheless, in the majority of these preparations (7/8), B63 was still able to produce plateau potentials in response to intracellular depolarizing current injection (Figure 4A2, D), thereby indicating that CPA did not in fact suppress this membrane property. In contrast, the non-selective cation channel blocker FFA (0.1 mM in a Low Ca+Co solution, n = 8) did not suppress B63’s spontaneous membrane oscillation (Figure 4B1, B2, C), which led to action potential bursts, but solely with short durations, on nearly every cycle. Moreover, in almost all of these tested preparations (7/8), experimental depolarization by current pulse injection, also failed to trigger any sustained plateau-like potentials (Figure 4B2, D). Quantitative comparisons between the amplitude of B63’s spontaneous oscillations and of the number of preparations generating plateau potentials that outlast a triggering stimulus pulse, thus indicated that these membrane properties are oppositely affected by FFA and CPA (Figure 4C, D). Therefore, although the B63 neuron’s oscillatory and plateauing capability both depend on TTX-sensitive membrane channels, the above pharmacological results indicated the involvement of two distinct types of channels.

**Figure 4.**
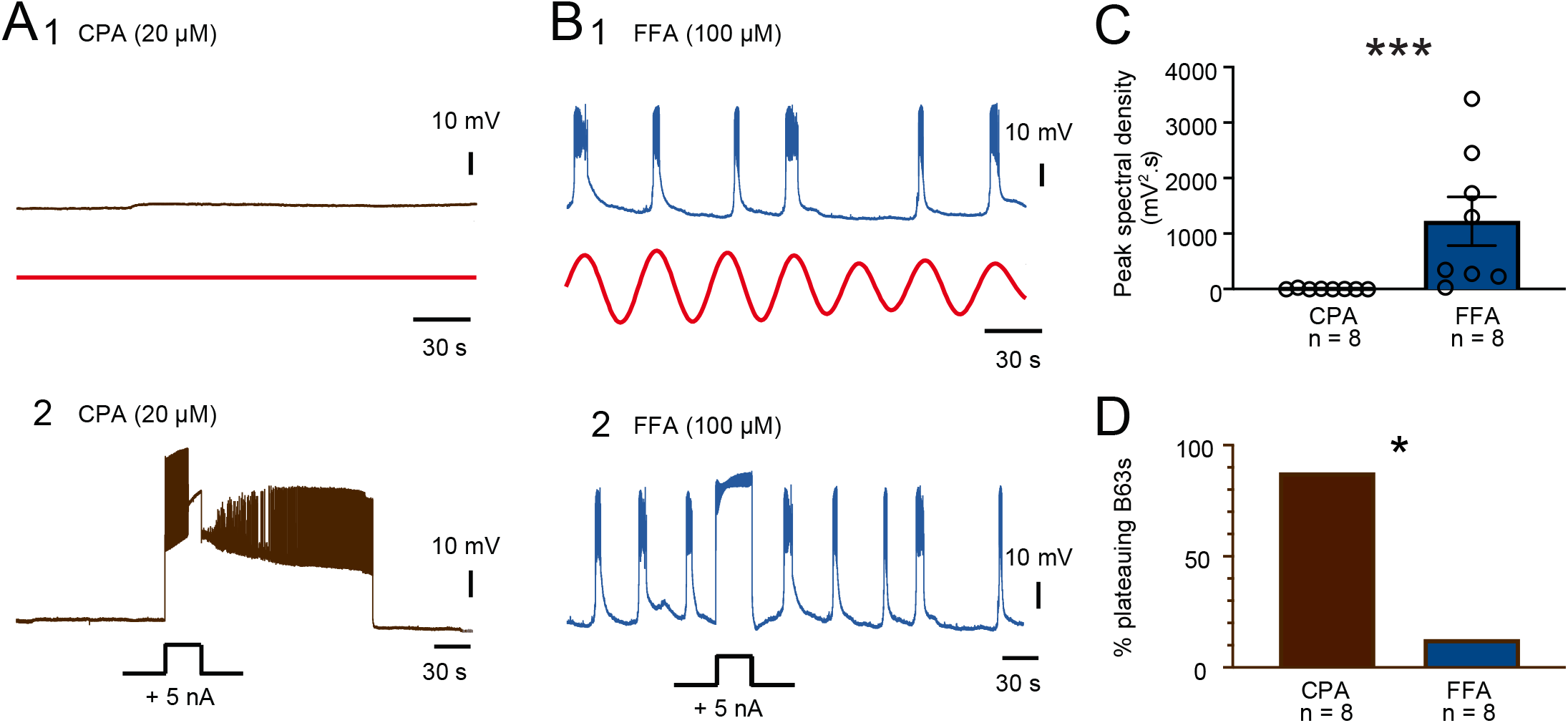
Differential sensitivity of B63 plateauing and oscillation to CPA and FFA. **A.** Effect of CPA. The presence of the SERCA blocker (20 μM in Low Ca+Co) suppressed a recorded B63’s spontaneous oscillation in membrane potential (A1, black trace, raw recording; red trace, reconstructed waveform from the peak FFT spectral density) but not the ability of an injected positive current pulse (+5 nA) to trigger a plateau potential (A2). **B.** Effect of FFA. The presence of the non-specific cation channel blocker (0.1 mM in Low Ca+Co saline) did not suppress expression of spontaneous membrane potential oscillation and associated short-lasting bursting activity in a B63 (B1, red trace: reconstructed waveform from the peak FFT spectral density), but it prevented expression of spontaneous or current pulse-evoked, long-lasting plateau potentials (B2, black trace: injected current). **C.** The amplitude of the spontaneous rhythmic oscillation in B63 (as see figure 1B, C) was significantly reduced in Low Ca+Co saline containing CPA as compared to FFA (*U* = 0, p < 0.001). **D.** The percentage of preparations expressing current pulse-evoked plateau potentials in B63 was significantly reduced in Low Ca+Co containing FFA as compared to CPA (p =0.010).

## DISCUSSION

The genesis of impulse bursting in central neuronal networks plays a fundamental role in decision-making processes and motor pattern genesis. In *Aplysia*, the decision-making leading to buccal motor cycle production depends on autonomous depolarizations and associated burst activity generated by several network neurons (including B63, B31/B32) that are interconnected by their electrical and chemical synapses (Susswein et al., 2002; Saada et al., 2009; Saada-Madar et al., 2012). In a two-neuron process, synapse-driven depolarization of the B31/B32 motoneurons by the B63 pacemaker neuron become long-lasting, thought to be due to voltage-activated muscarinic autapses that produce a plateau-like activation of these cells, which in turn provide sustaining feedback excitation to B63. The present study extends on this synapse-mediated process by showing that the B63 neuron can express plateau potential generation in the absence of B31/B32 activity. Thus, in addition to an underlying synaptic mechanism, the production of plateau-like potentials in the B63/B31/B32 subset is also likely to result, at least in part, from an endogenous membrane property of B63 itself. This is especially relevant since this pacemaker cell type also possesses an endogenous oscillatory property that can spontaneously initiate plateau potential production (see Figure 1; Bédécarrats et al., 2021). Thus, the decision-making process triggering each radula bite cycle in *Aplysia* evidently relies on a complex interplay between different intrinsic membrane properties and reciprocal electrical and chemical synapses amongst a subset of buccal network neurons.

Neuronal endogenous plateau properties derive from ionic transmembrane channels mediating persistent inward currents that can be triggered by brief synaptic inputs or spontaneous variations in membrane potential (Russell and Hartline 1978). The onset and maintenance of these plateau potentials have been previously found to depend on voltage-dependent L-type calcium channels (Svirskis and Hounsgaard, 1997), voltage-dependent sodium channels responsible for *I*_NaP_ (Crill, 1996; Elson and Selverston 1997), non-voltage dependent, calcium activated non-specific cationic channels (CAN) (Partridge and Swandulla, 1988; Zhang et al. 1995; Morisset and Nagy 1999) or an interplay between these cationic currents (Bouhadfane et al., 2013). B63’s spontaneous membrane potential oscillation that triggers plateau potentials was previously found to arise from an intracellular dynamic involving organelle calcium release (Bédécarrats et al., 2021). The present study now shows that these two intrinsic capabilities rely on different sets of conductances, with distinct electrical and pharmacological characteristics. Although both properties are TTX-sensitive, B63’s oscillation depends on a voltage-independent mechanism that is blocked by a reticulum calcium pump inhibitor (CPA) and is insensitive to the CAN channel blocker FFA (Bédécarrats et al., 2021). In contrast, B63’s plateauing ability which depends on active, voltage-dependent rising and falling phases, can be triggered by a transient depolarization and hyperpolarization respectively, and is not sensitive to CPA but is so to FFA. It is noteworthy that under FFA, however, the rising phase of each plateau is preserved, resulting in brief impulse burst firing, but a prolonged (lasting for 10s of secs to mins) depolarization (i.e., a plateau) and accompanying sustained discharge now fails to occur. This is therefore consistent with the likelihood that different ion channels are implicated in the onset and maintenance phases of the plateau. While voltage-dependent channels insensitive to FFA and conveying a persistent Na^+^ current (*I*_NaP_) could contribute to the rising phase, a CAN current (I_CAN_) that is sensitive to FFA may contribute to plateau maintenance. This in turn raises a possible contribution by intracellular calcium stores (the plateau persists in the presence of CPA, which increases the intracellular calcium concentration) or a voltage-dependent process. Such a mechanism was previously described for non-selective cationic channels contributing to stimulus afterdischarge in *Aplysia* bag cells (Wilson et al., 1996). Finally, plateau termination in B63 was found to be sensitive to transient membrane hyperpolarization, the K^+^channel blocker TEA, and to Ca^2+^ removal, all consistent with the involvement of a voltage- and calcium-activated potassium channel (Golowasch and Marder, 1992; Russo and Hounsgaard, 1996; Brezina and Weiss, 1995). Further characterization of these conductances and their interaction with those underlying B63’s oscillatory capability will thereby increase our understanding of the autonomous decision-making process for goal-directed actions in *Aplysia*.

## AUTHOR CONTRIBUTIONS

AB, RN designed and performed the experiments. AB performed the data analysis, made the initial version of the figures and wrote the first draft of the manuscript. JS, RN prepared and edited the final version of the manuscript. All authors contributed to the article and approved the submitted version.

## FUNDING

This research was funded by the French ‘Agence National de la Recherche’ with grant ANR-10-Idex-03–02 (to AB).

## Notes

### Competing Interest Statement

The authors have declared no competing interest.

